# MmuPV1 and HPV E6 Elevate Nuclear Accumulation of IKKɑ to Impair NF-κB Signaling to Promote Persistent Infection

**DOI:** 10.64898/2026.11.05.764775

**Authors:** Preston T Shepherd, Kayla R Duxbury, Zachary Morris, James C Romero-Masters

## Abstract

The mouse papillomavirus (MmuPV1) E6 oncoprotein is essential for persistent infection and cancer development. Previous studies identified amino acid R130 of MmuPV1 E6 as critical for its interaction with the tumor suppressor MAML1. Mutation of this residue (R130A) disrupts MAML1 binding and prevents disease development in mice. In this study, we demonstrate that MmuPV1 E6 also disrupts NF-κB signaling through interactions with proteins in the IκB kinase (IKK) complex. Specifically, the MmuPV1 E6^R130A^ mutant is defective in binding IKK complex proteins and fails to impair NF-κB signaling in mouse keratinocytes, suggesting that these interactions are necessary for E6-mediated disruption of this pathway. Furthermore, MmuPV1 E6 expression is associated with increased nuclear localization of IKKɑ, whereas cells expressing the E6^R130A^ mutant exhibit no such increase. These findings suggest that E6-mediated nuclear localization of IKKɑ depends on its interaction with IKK complex proteins. Importantly, we demonstrate that the human papillomavirus HPV16 E6 oncoprotein similarly impairs NF-κB signaling and increases nuclear IKKɑ localization in human keratinocytes, suggesting a shared mechanism between mouse and human papillomaviruses. Additionally, analysis of previously published RNA sequencing data reveals that MmuPV1 E6 induces several transcriptional changes associated with increased nuclear IKKɑ. Together, these findings demonstrate that both MmuPV1 and HPV16 E6 disrupt NF-κB signaling in association with increased nuclear IKKɑ localization. This study provides the first evidence of an E6 oncoprotein interacting with and changing subcellular localization of IKKɑ to disrupt cellular homeostasis.

**Importance:** In this study, we demonstrate that the mouse papillomavirus (MmuPV1) E6 oncoprotein interacts with proteins of the inhibitor of κB kinase (IKK) complex to impair NF-κB signaling, a key cellular pathway involved in regulating processes such as the innate immune response. We find that the MmuPV1 E6^R130A^ mutant exhibits reduced binding to IKK complex proteins and a diminished ability to impair NF-κB signaling, highlighting the importance of these interactions. Notably, we demonstrate that HPV16 E6 similarly impairs NF-κB signaling in human keratinocytes, revealing a shared biological function between mouse and human papillomavirus E6 proteins. Furthermore, both MmuPV1 and HPV16 E6 promote increased nuclear localization of IKKɑ, which may contribute to disrupted NF-κB signaling. Analysis of previously published RNA sequencing data also reveals transcriptional changes associated with increased nuclear IKKɑ. Together, these findings identify a previously unrecognized interaction between E6 and IKKɑ that may contribute to disrupting cellular homeostasis.

## Introduction

Papillomaviruses (PVs) are a large family of nonenveloped, small double-stranded DNA viruses that infect stratified squamous epithelia (1, 2). Persistent human papillomavirus (HPV) infections are responsible for ∼5% of the world’s total cancer burden (3–6). HPVs can be categorized into two categories based on tissue tropism - mucosal for anogenital infections and cutaneous for skin infections. HPV types that are linked to cancer are known as high-risk HPVs (2, 3, 5–7). Low-risk and cutaneous HPVs commonly cause genital and skin warts, respectively (7–10). A subset of cutaneous HPVs are associated with non-melanoma skin cancer in long-term immune- suppressed transplant patients and patients with the genetic disorder epidermodysplasia verruciformis (11, 12).

Nuclear factor kappa B (NF-κB) signaling is a multifunctional signaling cascade that regulates various cellular processes including proliferation, apoptosis (both pro- and anti- apoptotic pathways), angiogenesis, and the innate immune response (13). NF-κB is broken up into canonical and non-canonical NF-κB signaling (13, 14). A key regulatory complex in NF-κB signaling is the inhibitor of kappa B kinase – IKK – complex. The IKK complex is composed of 3 subunits: two catalytic subunits IKKɑ and IKKβ and the regulatory subunit IKKγ (NEMO) (15, 16). For canonical NF-κB signaling, the IKK complex promotes phosphorylation and degradation of IκB which releases hetero- and homodimers of NFkB1, RelA (p65), and c-Rel for nuclear translocation (13, 15). For non-canonical NF-κB signaling, IKKɑ promotes phosphorylation and proteasomal processing of NFkB2 which promotes nuclear translocation of p52/RelB heterodimers (14, 17, 18). One of the challenges in studying NF-κB signaling is its complexity as it is both anti-viral by activating the innate immune response and pro-tumorigenic by promoting cancer-associated inflammation, angiogenesis, and cell survival (13). Because of this complexity, the contribution of NF-κB signaling to HPV persistent infection remains unclear (19).

The HPV E6 and E7 oncogenes are drivers of persistent infection and promote carcinogenesis by altering host regulatory signaling cascades through direct interactions with key regulatory proteins within the cascade. The high-risk HPV E6 oncoprotein interacts with the LXXLL motif-containing E3 ubiquitin ligase E6AP to promote p53 degradation to circumvent the anti-proliferative activity of p53 (20–23). In contrast, cutaneous HPV E6 oncoproteins preferentially associate with the LXXLL motif-containing protein Mastermind-like 1 (MAML1) rather than E6AP (24–26). However, both high-risk and cutaneous HPV E6 oncoproteins interact with various other cellular proteins which likely contribute to persistent infection (27–30). These various activities highlight the complexity of the HPV E6 oncoprotein and the various activities that may promote persistent infection.

NF-κB signaling has been shown to be enriched in cervical cancer models (31). However, the impact of high-risk HPV E6 and E7 on NF-κB signaling varies based on the model system and cell line used (19, 28, 32–45). In non-cancerous cell models, it has been shown that high-risk HPV E6 and E7 impairs NF-κB signaling and NF-κB is antiproliferative in normal cervical cells (37, 38). HPV16 E7 has been shown to interact with and impair the IKK complex in overexpression models (37). Additionally, high-risk HPV E6 has been shown to block NF-κB signaling using overexpression models (35). In contrast, some studies have found that HPV16 E6 activates NF- κB signaling (28, 32, 39–41). Cutaneous HPVs also have been shown to impair NF-κB signaling in keratinocytes (42, 45, 46). In HPV positive cancer cells, NF-κB signaling is increased, suggesting a potential delineation between HPV’s impact on NF-κB signaling in normal cells versus cancer cells (31, 47). However, the exact mechanisms by which various HPVs impact NF- κB signaling in persistent infection models remain unresolved.

While the IKK complex has a well-defined role in regulating NF-κB signaling, IKKɑ has been shown to have NF-κB-independent activities (48–51). IKKɑ has been shown to regulate skin development, differentiation, and keratinocyte proliferation (48–54). IKKɑ has been identified as a tumor suppressor, and mice with conditional IKKa knockout are more susceptible to squamous cell carcinoma development (48–54). A recent study examining the impact of nuclear versus cytoplasmic IKKɑ found distinct and opposing roles of IKKɑ in skin biology (51). In the cytoplasm, IKKɑ is pro-inflammatory but also contributes to keratinocyte proliferation (51). In contrast, nuclear IKKɑ is anti-inflammatory and increases c-Myc activity and alters metabolism (51). Overexpression of IKKɑ in the nucleus did lead to spontaneous squamous cell carcinoma development in mice (51). The impact of HPV E6 or E7 oncoproteins on NF-κB independent activities of IKKɑ and IKKɑ localization has never been studied before.

Our ability to study the mechanisms by which HPV promotes persistent infection *in vivo* is not possible due to the species specificity of papillomaviruses. We study the murine papillomavirus MmuPV1 as a model system for persistent HPV infection (55, 56). Importantly, MmuPV1 infects and causes disease including cancer at clinically relevant sites (56–64). MmuPV1 E6 is required for persistent infection like HPV E6 (65). We recently published a study showing that MmuPV1 E6 has a significant impact on keratinocyte homeostasis by increasing proliferation, blocking p53 signaling, altering metabolism, and impairing the innate immune response (66). We found that a mutant MmuPV1 E6, E6^R130A^, was significantly impaired in promoting proliferation of mouse keratinocytes *in vitro* (66). The MmuPV1 E6^R130A^ mutant has been previously shown to be defective for promoting disease *in vivo* which was linked to a significant reduction in the interaction between MmuPV1 E6 and MAML1 (65, 67). MmuPV1 E6’s interaction with MAML1 contributes to MmuPV1 E6’s ability to impair differentiation (65, 67). However, MmuPV1 E6 has also been shown to interact with SMAD2/3 and this interaction is not impacted by the MmuPV1 E6^R130A^ mutant (65). MmuPV1 E6 likely has other interacting partners that may contribute to persistent infection including targeting NF-κB signaling.

In this study, we examine the impact of MmuPV1 E6 on NF-κB signaling in mouse keratinocytes. We found that MmuPV1 E6 interacts with all the components of the IKK complex and the MmuPV1 E6^R130A^ mutant is defective in this interaction. MmuPV1 E6’s interaction with the IKK complex is associated with a decrease in NF-κB signaling when MmuPV1 E6 is overexpressed and in the context of the intact viral genome. We also find that HPV16 E6 also impairs NF-κB signaling in the context of human keratinocytes. Importantly, we show that MmuPV1 and HPV E6 increases the amount of nuclear IKKɑ in keratinocytes. We also find that the increase in nuclear IKKɑ correlates with transcriptional changes observed in previous studies on nuclear IKKɑ (51). Thus, our study highlights a new activity of MmuPV1 E6 that is shared with the high-risk HPV16 E6 oncoprotein that likely contributes to persistent infection.

## Results

### MmuPV1 E6 reduces NF-κB signaling in mouse keratinocytes

Our prior work has established that MmuPV1 E6 disrupts several host signaling cascades, i.e., NF-κB, which leads to increasing cellular proliferation, inhibiting tumor suppressor signaling, and impairing innate immune signaling (66). Gene set enrichment analysis (GSEA) of our published RNA-seq data suggests that MmuPV1 E6 reduces NF-κB signaling in mouse keratinocytes (MKs) (66). Specifically, the Hallmark TNFɑ signaling via NF-κB gene set showed significant negative enrichment in MmuPV1 E6-expressing MKs compared to vector control MKs (**Fig. 1A**). We further assessed NF-κB signaling by performing immunoblot analysis for phosphorylation of the NF-κB subunit p65 (RelA) at Ser536 (P-p65) in MmuPV1 E6-expressing MKs compared to vector control MKs. We observed a reduction in P-p65 in the MmuPV1 E6-expressing MKs compared to vector control MKs, with no apparent difference in total p65 protein levels (**Fig. 1B & C**). To functionally assess impairment of NF-κB signaling, we performed qRT-PCR for the NF-κB-responsive gene IL-6. We observed a significant reduction in IL-6 in MmuPV1 E6-expressing MKs (**Fig. 1D**). These data suggest that MmuPV1 E6 reduces NF-κB signaling in mouse keratinocytes.

**Figure 1.**
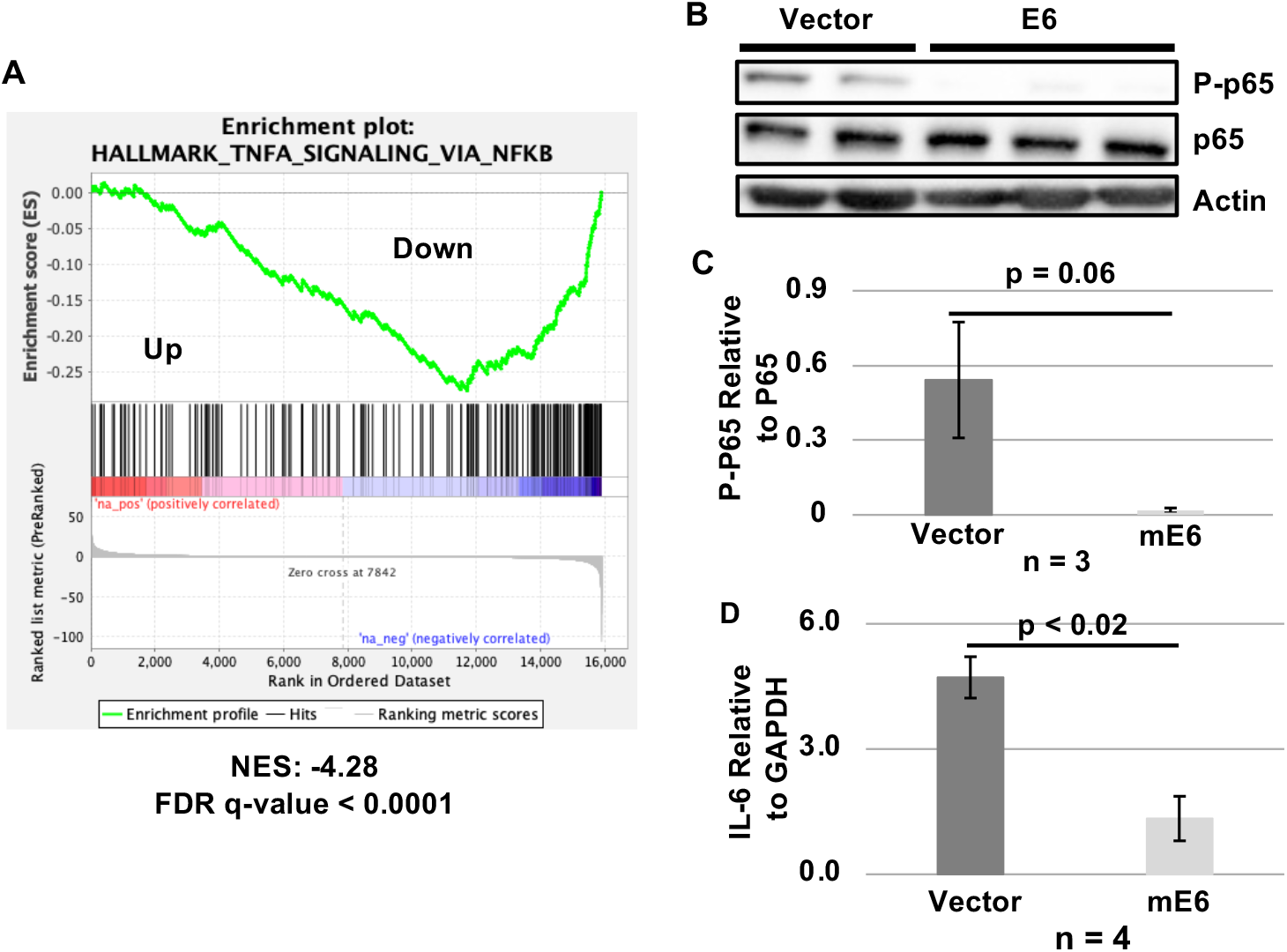
MmuPV1 E6 reduces NF-κB signaling in mouse keratinocytes. **(A)** GSEA of published RNA-seq data comparing MmuPV1 E6-expressing and vector control mouse keratinocytes (MKs) (66). The Hallmark_TNFA_Signaling_via_NF-kB gene set is shown. We observed a significant negative enriched in MmuPV1 E6-expressing MKs compred to vector control MKs. **(B)** Protein extracts from MmuPV1 E6-expressing and vector control MKs were subject to immunoblot analysis using antibodies against phosphorylated p65 (P-p65) at serine 536 – a known IKK complex phosphorylation site, total p65, and actin. **(C)** Western blot quantification of panel B with a reduction in P-p65 levels. Quantification was performed using ImageJ. Vector was in biological duplicate (n=2) and MmuPV1 E6-expressing MKs was in biological triplicate (n=3). **(D)** qRT-PCR was performed on cDNA generated from RNA isolated from MmuPV1 E6-expressing and vector control MKs using primers targeting the mouse IL-6 gene. Pair-wise t-tests were performed and standard error is shown. Experiment was performed in biological quadruplicate (n=4).

### MmuPV1 E6^R130A^ does not increase the proliferation rate of mouse keratinocytes *in vitro*

We have previously shown that MmuPV1 E6 increases the proliferation capacity of mouse keratinocytes (MKs) *in vitro* and the MmuPV1 E6^R130A^ mutant is impaired in promoting proliferation of MKs (66). This reduction in proliferation correlates with the E6^R130A^ being unable to cause papillomas in mice (65). However, our prior studies used an N-terminal HA-tagged MmuPV1 E6 which may impair some of MmuPV1 E6’s activities. To further solidify this observation, we repeated our growth analysis comparing wild-type (WT) MmuPV1 E6-expressing, E6^R130A^- expressing, and vector control MKs using an untagged retrovirus expression construct. We generated WT MmuPV1 E6-expressing, E6^R130A^-expressing, and vector control MKs and confirmed E6 expression using qRT-PCR (**Fig 2A**). These MK cells strains were then subjected to our published growth analyses (66). We found that MmuPV1 E6^R130A^-expressing MKs showed reduced proliferation compared with WT MmuPV1 E6-expressing MKs in both assays (**Fig. 2B & C**). The MmuPV1 E6^R130A^-expressing MKs proliferated at a rate similar to that of vector control MKs in our short-term growth assay (**Fig. 2B**). We observed a significant reduction in the total fold increase in MmuPV1 E6^R130A^-expressing MKs compared to WT MmuPV1 E6-expressing MKs in our long-term assay as expected (**Fig. 2C**). Interestingly, we did observe a separation between the MmuPV1 E6^R130A^-expressing MKs and vector control MKs in the long-term growth assay (**Fig. 2C**), suggesting that the mutant retains a slight degree of pro-proliferative activity. Collectively, our observations confirm that the interactions facilitated by MmuPV1 E6 through amino acid R130 contribute to MmuPV1 E6’s pro-proliferative activity.

**Figure 2.**
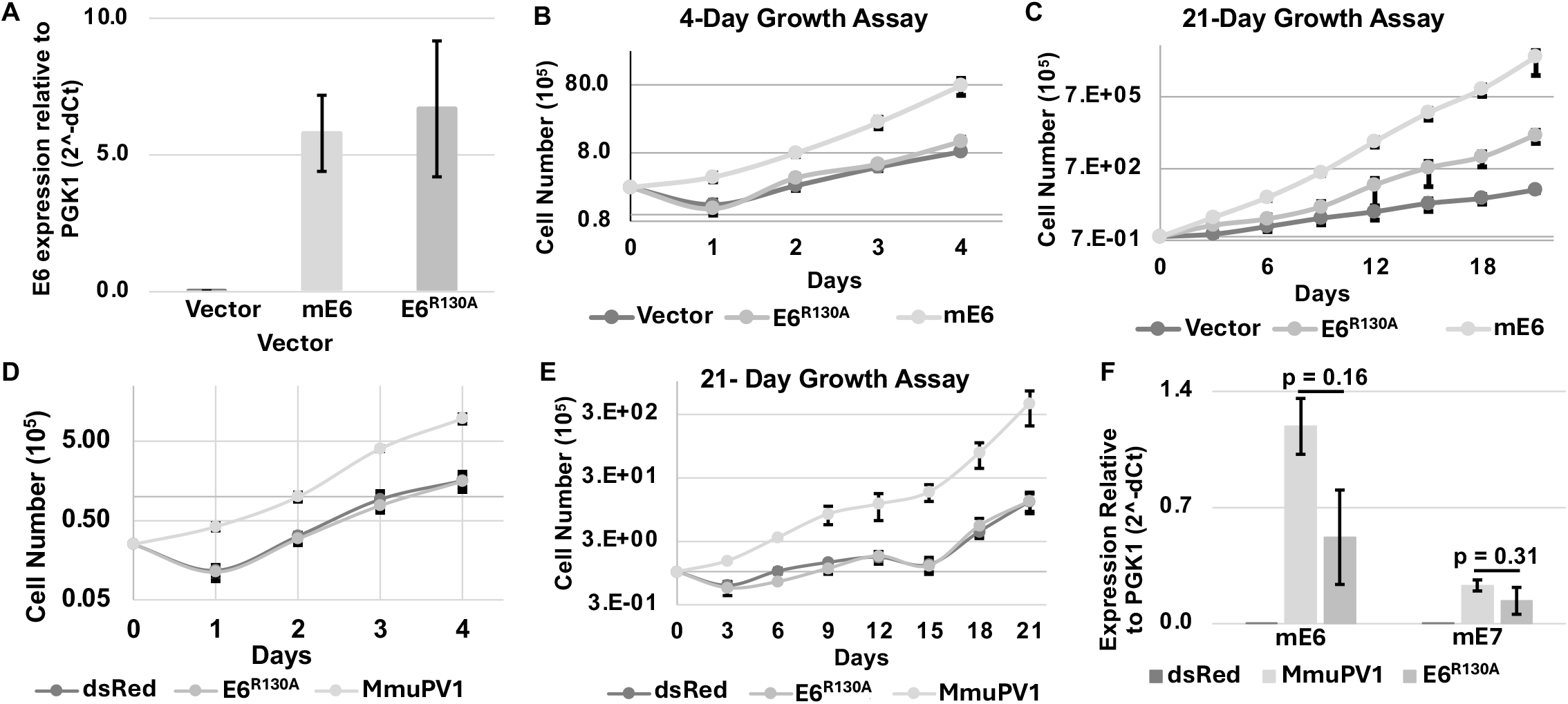
MmuPV1 E6^R130A^ is defective in promoting proliferation. (**A)** qRT-PCR analysis was performed on cDNA generated from RNA isolated from vector control, MmuPV1 E6- (mE6), and E6^R130A^ -expressing mouse keratinocytes (MKs). We observe no significant difference in E6 gene expression in our MKs. **(B**) 4-day growth analysis comparing MmuPV1 E6- (mE6), E6^R130A^-, and vector control MKs. We observed a significant increase in growth in MmuPV1 E6-expressing MKs and the E6^R130A^-expressing MKs grew at a rate similar to vector control MKs. Linear regression analysis was performed to determine significance – E6 vs Vector: p < 0.02, E6 vs E6^R130A^: p < 0.05, and E6^R130A^ vs Vector: p = 0.75. (**C**) 21-day growth assay was performed comparing MmuPV1 E6- (mE6), E6^R130A^-, and vector control MKs. There was a significant increase in the cumulative fold increase in MmuPV1 E6-expressing MKs compared to E6^R130A^-expressing (p < 0.001) and vector control MKs (p < 0.001). MmuPV1 E6^R130A^-expressing MKs had a significant increase in growth compared to vector control (p < 0.001). (**D**) 4-day growth assay of MKs harboring recircularized full-length WT MmuPV1, MmuPV1 E6^R130A^ genomes, and dsRed control MKs. WT MmuPV1 increased the proliferation rate of MKs compared to both dsRed control (p < 0.04) and E6^R130A^ (p < 0.05). **(E)** 21-day growth assay of MKs harboring recircularized full-length WT MmuPV1, MmuPV1 E6^R130A^ genomes, and dsRed control MKs. WT MmuPV1 significantly increased the cumulative growth rate of the MKs compared to dsRed control (p < 0.001) and E6^R130A^ (p < 0.002). **(F)** qRT-PCR was performed for MmuPV1 E6 and E7 on cDNA generated from RNA isolated from WT MmuPV1, E6^R130A^ mutant, and dsRed control MKs. 2^−ΔCt^ values are plotted. No significant difference in E6 or E7 expression was detected. Standard error is shown in all panels. All experiments were performed in biological quadruplicate (n=4).

To better establish the relevance of MmuPV1 E6’s ability to increase proliferation in MKs during an MmuPV1 infection, we generated MK cell strains that maintain the intact MmuPV1 genome. To do this, we transfected MKs with dsRed (encodes a Blasticidin resistance gene for selection) alone or in combination with recircularized WT MmuPV1 or MmuPV1 E6^R130A^ genomes, as previously published (68). We confirmed the presence of the MmuPV1 genome and subjected these cells to our growth analyses. In both growth assays, WT MmuPV1 increased MK proliferation compared to the dsRed control cells (**Fig 2D & E**). The MmuPV1 E6^R130A^ genome containing MKs showed a proliferation rate comparable to the dsRed MKs and significantly lower than WT MmuPV1 genome containing MKs in both assays (**Fig. 2D & E**). To confirm that this reduction in growth was not due to differences in the transcription of MmuPV1 E6 or E7, we performed qRT-PCR on cDNA generated from RNA isolated from our MKs. We did not observe a significant difference in the transcription of MmuPV1 E6 or E7 in our analysis (**Fig. 2F**). Collectively, our observations show that MmuPV1 E6’s ability to interact with LXXLL-motif containing proteins like MAML1 contribute to the increase in proliferation of mouse keratinocytes *in vitro*.

### MmuPV1 E6 interacts with the IKK complex

To investigate how MmuPV1 E6 impairs NF-κB signaling, we wanted to determine whether MmuPV1 E6 directly interacts with NF-κB signaling proteins. A prior interactomic study (affinity-purification mass spectrometry or AP/MS), performed by the Munger lab, found that MmuPV1 E6 pulled down peptides from various proteins in the NF- κB signaling cascade including IKKɑ, IKKβ, IKKγ, NFkB1, NFkB2, and IκBɑ (65). We performed coimmunoprecipitation-western blot analysis (Co-IP/WB) on transiently transfected HCT116 cells which were used in the prior interactomic study (65). MAML1, IRF3, and SMAD2/3 co- immunoprecipitated with WT MmuPV1 E6 as expected (**Fig. 3A**). We confirmed that MmuPV1 E6 interacts with all the NF-κB signaling proteins that were identified in the prior AP/MS analysis (**Fig 3A**). E6AP did not co-immunoprecipitate with WT MmuPV1 E6, as expected, and acted as a negative control for the Co-IP/WB analysis (**Fig. 3A**). Unexpectedly, we found that the MmuPV1 E6^R130A^ did not interact with any of the NF-κB signaling proteins including IKKɑ, IKKβ, IKKγ, NFkB1, and NFkB2 (**Fig. 3A**). Therefore, our data suggests that MmuPV1 E6 interacts with various members of the NF-κB signaling cascade.

**Figure 3.**
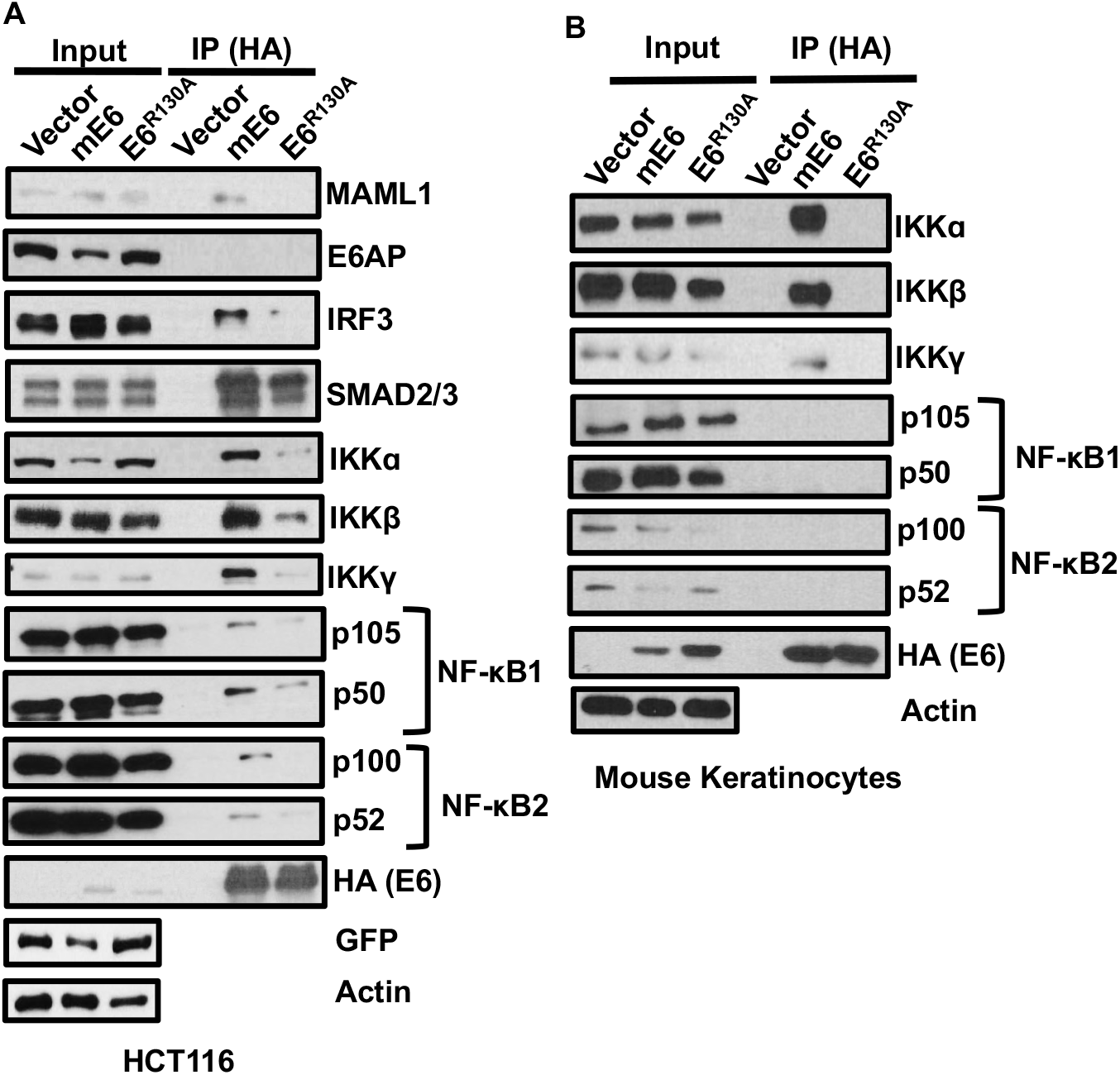
MmuPV1 E6^R130A^ mutant is defective for interacting with the IKK complex. (**A**) Co- immunoprecipitation/western blot (CoIP/WB) analysis was performed on HCT116 cells transiently transfected with vector control, HA-tagged wild-type MmuPV1 E6 (mE6), or HA-tagged MmuPV1 E6^R130A^. Immunoblot analysis was performed on 10 μg inputs and immunoprecipitants using antibodies against MAML1, E6AP (UBE3A), IRF3, SMAD2/3, IKKɑ, IKKβ, IKKγ, NFkB1, NFkB2, HA (detects MmuPV1 E6), GFP, and actin. Representative blot is shown. Experimental triplicate (n = 3) was performed for analysis. (**B**) CoIP/WB analysis was performed on mouse keratinocytes stably expressing vector control, HA-tagged WT E6 (mE6), or HA-tagged E6^R130A^. Immunoblot analysis was performed on 10 μg inputs and immunoprecipitants using antibodies against IKKɑ, IKKβ, IKKγ, NFkB1, NFkB2, HA (detects MmuPV1 E6), and actin. Representative blots are shown from biological quadruplicate (n = 4) experiments.

To validate our results in a physiologic relevant cellular environment, we generated mouse keratinocytes (MKs) that stably expressed HA-tagged WT MmuPV1 E6, HA-tagged MmuPV1 E6^R013A^, and vector control using a retroviral expression construct. Following selection and confirmation of E6 expression, we subjected these MKs to Co-IP/WB analysis. Consistent with our HCT116 cell experiments, WT MmuPV1 E6 interacted with IKKα, IKKβ, and IKKγ in MKs, whereas no detectable association was observed with NF-κB1 or NF-κB2 (**Fig. 3B**). The MmuPV1 E6^R130A^ mutant was defective for interacting with IKKɑ, IKKβ, and IKKγ in MKs (**Fig. 3B**). Collectively, our observations suggest that MmuPV1 E6 interacts with IKKɑ, IKKβ, and IKKγ and the MmuPV1 E6^R130A^ mutant is defective for these interactions.

### MmuPV1 E6 impairs NF-κB signaling through its interaction with the IKK complex

We next examined the impact of MmuPV1 E6’s interaction with IKKα, IKKβ, and IKKγ on NF-κB signaling in mouse keratinocytes (MKs). As shown in our initial studies, MmuPV1 E6 reduces phosphorylation of p65 (RelA) at S536 a known IKK complex phosphorylation site (**Fig. 1B**) (69). To more broadly assess the impact of MmuPV1 E6 on NF-κB signaling, we performed immunoblot analysis for NFkB2 on protein extracts isolated from MKs that stably expressed WT MmuPV1 E6 or MmuPV1 E6^R130A^, and vector control MKs. We found that MmuPV1 E6 increased the abundance of the unprocessed p100 relative to p52 which is an IKKɑ regulated process (**Fig. 4A & B**). The MmuPV1 E6^R130A^-expressing MKs showed a pattern similar to vector controls (**Fig. 4A & B**). Consistent with what we observed in **Fig 1**, WT MmuPV1 E6-expressing MKs showed reduced p65 Ser536 phosphorylation compared with vector control (**Fig. 4C & D**). The MmuPV1 E6^R130A^-expressing MKs showed a phosphorylation pattern similar to vector control MKs (**Fig. 4C & D**). These data suggest to us that MmuPV1 E6 overexpression reduces baseline NF-κB signaling.

**Figure 4.**
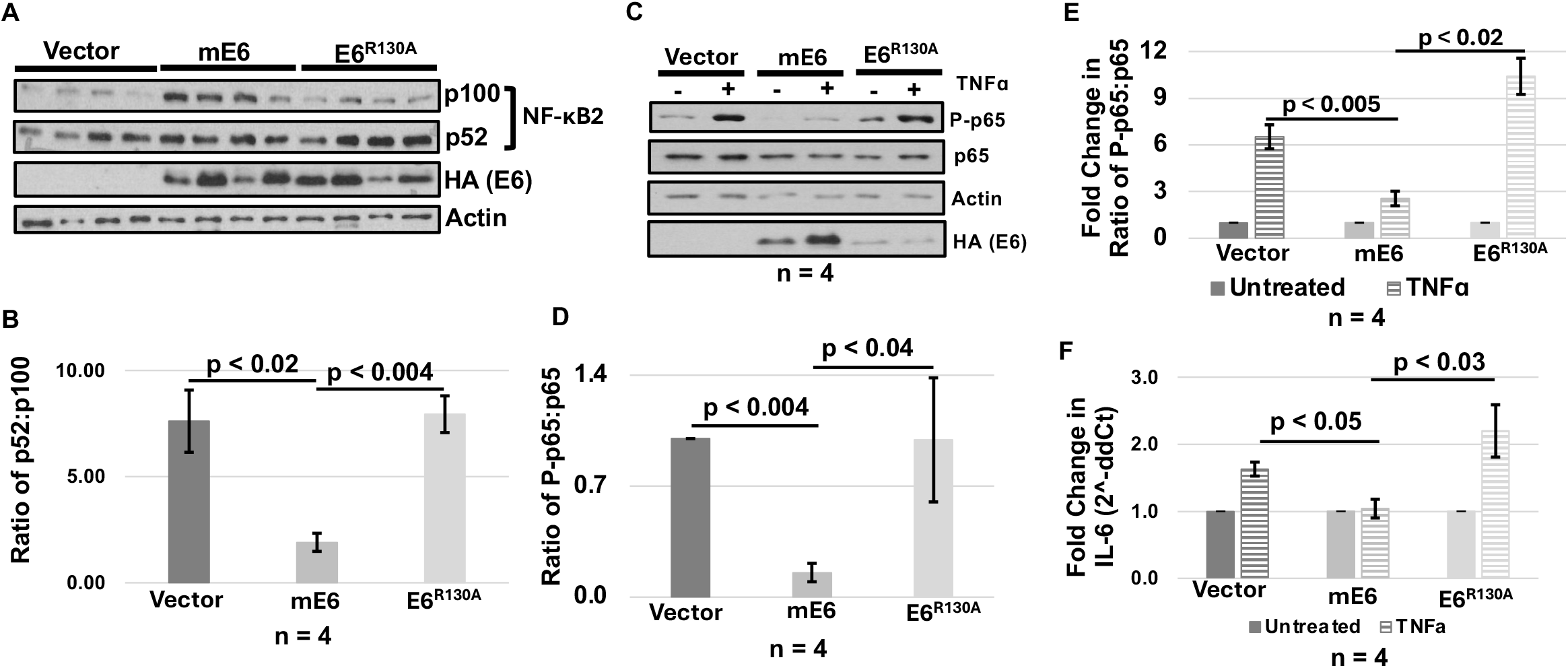
MmuPV1 E6’s interaction with IKK complex contributes to impairment of NF-κB signaling in mouse keratinocytes. (**A**) Immunoblot analysis was performed on mouse keratinocytes (MKs) stably expressing vector control, HA-tagged wild-type (WT) MmuPV1 E6 (mE6), or HA-tagged MmuPV1 E6^R130A^ using antibodies against NFkB2, HA (detected MmuPV1 E6), and actin. (**B**) Quantification of NFkB2 immunoblot analysis in panel **A** is shown. Quantification compared the ratio of p52 (processed) and p100 (unprocessed) NFkB2. (**C**) Immunoblot analysis was performed on extracts isolated from stable-expressing MKs described above treated with 10 ng/mL TNFɑ (+) or untreated (-). Antibodies against Phospho-p65 (P-p65) (Ser536), total p65, actin, and HA (detects MmuPV1 E6) were used in immunoblot analysis. Representative blot is shown. Experiment was performed in biological quadruplicate (n=4). (**D**) Quantification of P-p65 relative to total p65 in untreated conditions in panel **C** is shown. Experiment was performed in biological quadruplicate (n=4). Pair-wise t-test was performed. (**E**) Quantification of immunoblot analysis shown in panel **C** comparing the fold increase of P-p65 when treated with TNF-ɑ. Fold-change was relative to untreated condition. Experiment was performed in biological quadruplicate (n=4). Pair-wise t-test was performed. **(F)** qRT-PCR was performed using primers for IL-6 and PGK1 on cDNA generated from RNA isolated from vector control, WT MmuPV1 E6-, or MmuPV1 E6^R130A^- expressing MKs. Fold change in IL-6 was determined comparing treated to untreated condition. 2^-ΔΔCt^ was plotted and is shown. Experiment was performed in biological quadruplicate. Pair-wise t-test was performed. All graphs show standard error.

Considering NF-κB responds to various stimuli, we wanted to determine whether MmuPV1 E6 reduces the ability of MKs to respond to TNFɑ stimulation comparing our WT MmuPV1 E6- expressing, MmuPV1 E6^R130A^-expressing, and vector control MKs. Following plating, cells were grown to near confluency and then stimulated with 10 ng/mL of TNFɑ. We found that vector control MKs showed an increase in phosphorylated p65 at Ser536 (**Fig. 4C & E**). In contrast, WT MmuPV1 E6-expressing MKs showed a significantly lower increase in phosphorylation of p65 at Ser536 than vector control (**Fig. 4C & E**). Importantly, MmuPV1 E6^R130A^-expressing MKs showed an induction similar to vector control MKs and was significantly higher than WT MmuPV1 E6- expressing MKs (**Fig. 4C & E**). To assess the impact on transcriptional changes associated with NF-κB signaling, we performed qRT-PCR analysis for the NF-κB responsive gene IL-6 after TNFɑ stimulation. Consistent with our immunoblot analysis, we found that WT MmuPV1 E6-expressing MKs have lower IL-6 expression following TNF-ɑ stimulation than vector control MKs (**Fig. 4F**) Consistent with immunoblot analysis, MmuPV1 E6^R130A^-expressing MKs showed an increase in IL-6 expression comparable to vector control MKs and significantly higher than WT MmuPV1 E6- expressing MKs (**Fig. 4F**). Collectively, our results suggest MmuPV1 E6 impairs NF-κB signaling through its interaction with IKK complex proteins.

### MmuPV1 infection impairs NF-κB signaling *in vitro* and *in vivo*

We next wanted to translate the impact of MmuPV1 E6 on NF-κB signaling during an MmuPV1 infection. To do this, we generated mouse keratinocytes (MKs) that maintain the intact viral genome. The cells used in this analysis were generated in **Fig 2**. To determine if MmuPV1 E6 expression from the intact viral genome was sufficient to impair baseline NF-κB signaling, we performed immunoblot analysis of NF-κB2. We observe that MmuPV1 containing cells did not significantly change the amount of p100 protein present compared to p52 (**Fig. 5A & B**). Similar to what we observed in MmuPV1 E6 overexpression, WT MmuPV1 genome-containing MKs has significantly lower levels of phosphorylated p65 at S536 compared to dsRed control (**Fig. 5C & D**). Additionally, the MmuPV1 E6^R130A^ genome-containing MKs had similar levels of phosphorylated p65 as dsRed control MKs and a significantly higher levels than WT MmuPV1 MKs (**Fig. 5C & D**). To assess the impact of the intact viral genome on NF-κB following TNFα stimulation, we performed identical experiments as described in **Fig 4**. Similar to what we observed in MmuPV1 E6 overexpression, WT MmuPV1 genome-containing MKs had a significantly lower response to TNF-ɑ stimulation compared dsRed control MKs (**Fig. 5C & 5E**). As expected, the MmuPV1 E6^R130A^ genome containing MKs has a stimulation response similar to dsRed control MKs and a significantly higher response than WT MmuPV1 genome-containing MKs (**Fig. 5C & 5E**). These results suggest that MmuPV1 E6 impairs NF-κB signaling in the context of the intact viral genome and this impairment is dependent on the MmuPV1 E6’s interaction with the IKK complex.

**Figure 5.**
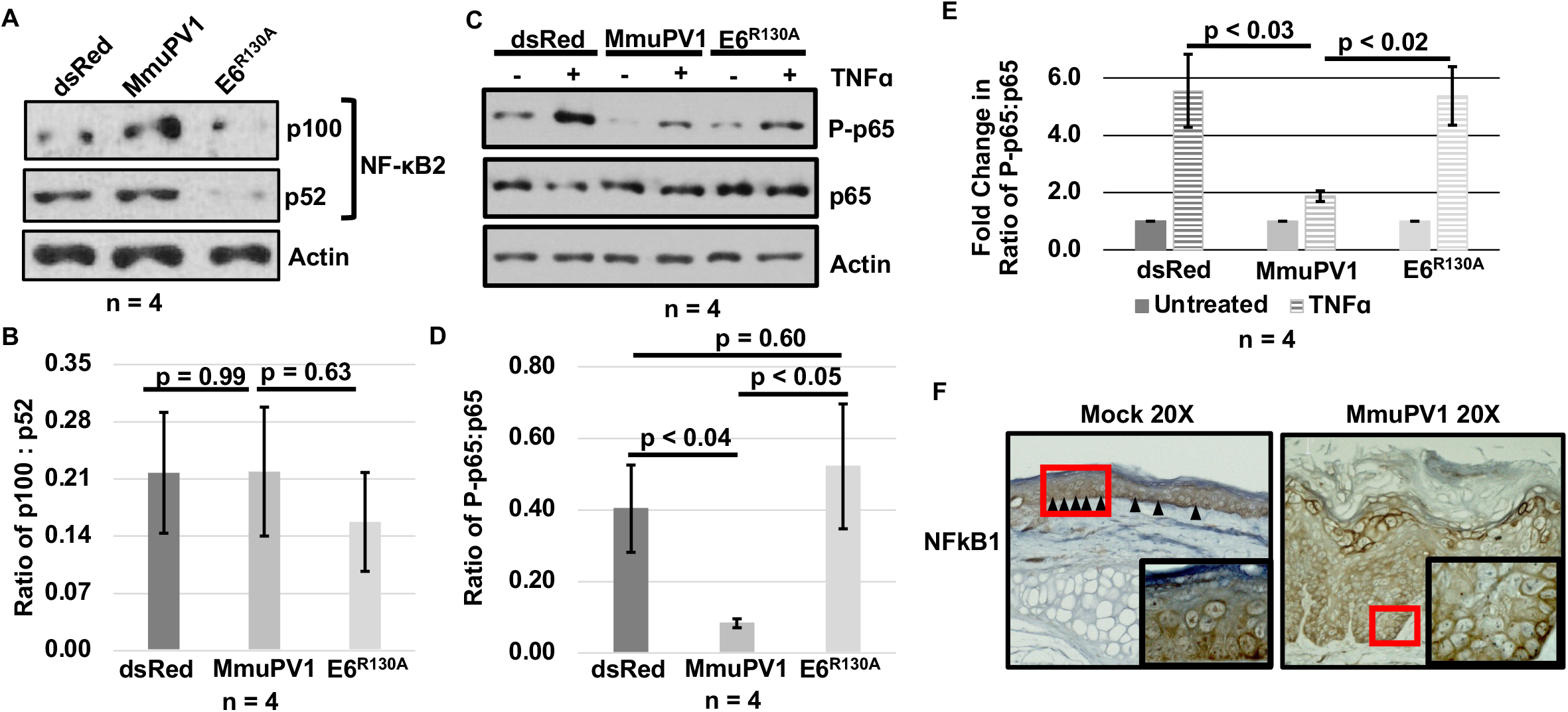
MmuPV1 infection attenuates NF-κB signaling both *in vitro* and *in vivo.* (**A)** Immunoblot analysis on extracts isolated from mouse keratinocytes (MKs) that maintain either wild-type (WT) MmuPV1 or MmuPV1 E6^R130A^ intact viral genome and dsRed control MKs. Immunoblots used antibodies against NFkB2 and actin. Representative blot is shown. Experiment was performed in biological quadruplicate. **(B**) Quantification of NF-κB2 immunoblot analysis shown in panel A. Bar graph shows the ratio of p100:p52. Standard error is shown and experiment was performed in biological quadruplicate (n=4). (**C)** Immunoblot analysis on intact viral genome containing MKs described above treated with 10 ng/mL TNF-ɑ (+) or untreated (-) for 10 minutes. Antibodies against Phosphorylated p65 S536 (P-p65), total p65, and actin were used in our immunoblot analysis. Representative blot is shown. Experiment was performed in biological quadruplicate (n=4). **(D)** Quantification of baseline P-p65 relative to total p65 of immunoblot analysis shown in panel **C**. Experiment was performed in biological quadruplicate (n=4). (**E**) Quantification of immunoblot analysis shown in panel **C** comparing the fold increase in P-p65 when treated with TNF-ɑ. Fold-change is relative to untreated condition. Experiment was performed in biological quadruplicate (n=4). Pair-wise t-test was performed in all bar graphs shown and standard error is show in all graphs. (**F**) Immunohistochemistry (IHC) was performed using antibodies against NFkB1. Both mock and MmuPV1-infected tissue sections were stained in our analysis. Red boxes mark the regions enlarged in the insets. Images were acquired with a 20X or 63X objective.

MmuPV1’s power as a model is in the ability to examine the role of MmuPV1 E6 during an infection *in vivo*. Considering our *in vitro* results, we wanted to determine the impact of MmuPV1 persistent infection on NF-κB signaling in an MmuPV1-infected lesions. To do this, we performed immunohistochemistry (IHC) for NFkB1. We examined our stained tissues for nuclear localization of NFkB1. We found that MmuPV1 infected lesions had an absence of nuclear NFkB1 staining (**Fig. 5F**). Considering NF-κB signaling is not constitutively active in normal skin, we were not surprised by the faint nuclear NFkB1 staining in our Mock infected tissues (**Fig. 5F**). Unfortunately, we are unable to compare WT MmuPV1 to E6^R130A^ due to the E6^R130A^ mutant being unable to cause disease *in vivo*. Our *in vivo* analysis suggests that MmuPV1 infection also reduces NF-κB signaling during a persistent infection *in vivo*.

### HPV16 E6 impairs TNFɑ-induced NF-κB signaling in human keratinocytes

The impact of high-risk HPV16 on NF-κB signaling varies depending on the model system (28, 32, 37, 38). We wanted to better address the impact of high-risk HPV16 E6 on NF-κB in normal human keratinocytes. To address this question, we used N/TERT1 immortalized normal human keratinocytes that either express HPV16 E6 (16E6), HPV16 E7 (16E7), or HPV16 E6E7 (16 E6E7) and control N/TERT1 cells. We included HPV16 E7 in our analysis as prior literature has shown an interaction between HPV16 E7 and IKK complex proteins (37). We first performed immunoblot analysis for NFkB1 and NFkB2 processing in 16E6, 16E7, and 16E6E7 N/TERT1 cells compared to N/TERT1 cells. We did not observe any change in baseline in processing of NFkB1 and NFkB2 (**Fig. 6A**). We also did not observe a significant impact on baseline P-p65 at S536 between any of our conditions (**Fig. 6B**). Therefore, we wanted to determine the impact of HPV16 E6 and E7 expression alone or in combination on TNFɑ stimulation. We treated our N/TERT1, 16E6-expressing, 16E7-expressing, and 16E6E7-expressing cells with 10 ng/mL TNF-ɑ. We initially did the experiment where N/TERT1 cells were grown in the absence of feeders which is the standard approach for growing N/TERT1 cells (**Fig 6B & C**). We found that P-p65 was strongly induced by TNF-ɑ when N/TERT1, 16E6, 16E7, and 16E6E7 cells were grown without feeders with no significant differences (**Fig. 6B & C**). Considering that our MmuPV1 E6 mouse keratinocyte (MK) experiments were done in the presence of feeders, we repeated the TNF-ɑ stimulation study in the presence of feeders. Using the same methods in our MK studies, we found that 16E6 and 16E6E7 significantly impaired phosphorylation of p65 following TNF-ɑ stimulation compared to N/TERT1 cells (**Fig. 6D & E**). This data suggests to us that 16E6 impairs NF-κB signaling following TNFɑ stimulation in human keratinocytes. We did not observe an impact on P-p65 when 16E7 was expressed alone (**Fig. 6D & E**). 16E6 and 16E7 expression was confirmed using p53 and pRB (**Fig. 6**). Collectively, these results suggest that HPV16 E6, like MmuPV1 E6, impairs NF-κB signaling in human keratinocytes when the interaction between keratinocytes and stromal cells (fibroblasts) is mimicked.

**Figure 6.**
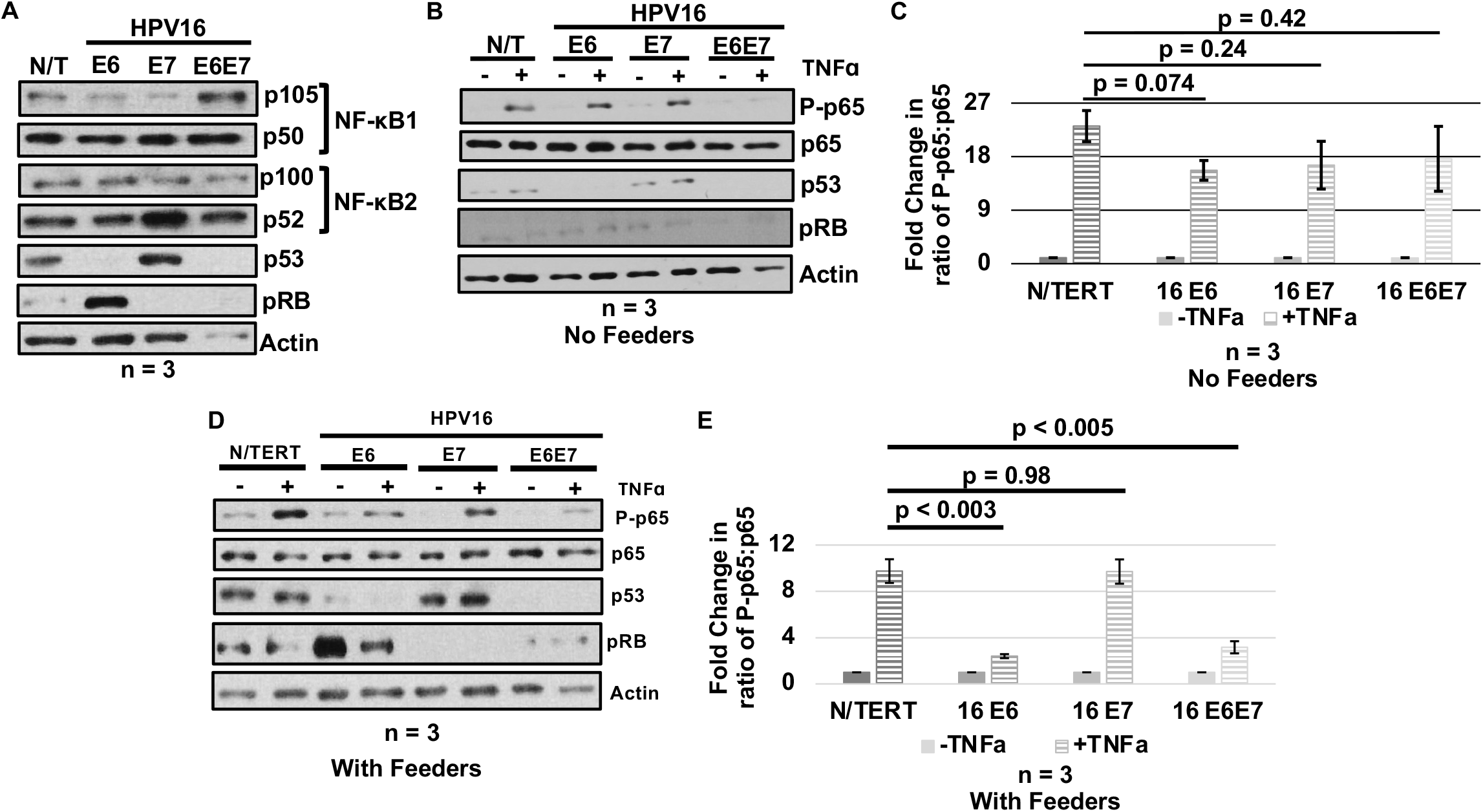
HPV16 E6 impairs TNFα-induced NF-κB signaling in human keratinocytes grown with feeders. **(A)** Immunoblot analysis was performed using antibodies against NFkB1, NFkB2, p53 (16E6 expression control), pRb (16E7 expression control), and actin on extracts from control, HPV16 E6- (E6), HPV16 E7- (E7), and HPV16 E6E7-expressing (E6E7) N/TERT1 (human keratinocytes) cells. Representative image is shown. **(B)** Immunoblot analysis was performed on extracts generated from control, HPV16 E6-, HPV16 E7-, and HPV16 E6E7-expressing N/TERT1 treated with 10 ng/mL TNF-ɑ (+) or untreated (-) when grown in the absence of feeders. Antibodies against Phosphorylated p65 at S536 (P-p65), total p65, p53, pRB, and actin were used in immunoblot analysis. **(C)** Quantification of immunoblot analysis shown in panel **B** comparing the fold increase in P-p65 when treated with TNF-ɑ. Fold-change is relative to untreated condition. **(D)** Immunoblot analysis was performed on extracts from control, HPV16 E6-, HPV16 E7-, and HPV16 E6E7-expressing N/TERT1 treated with 10 ng/mL TNF-ɑ (+) or untreated (-) when grown in the presence of feeders. Antibodies against Phosphorylated p65 at S536 (P-p65), total p65, p53, pRB, and actin were used in immunoblot analysis. **(E**) Quantification of immunoblot analysis shown in panel **D** comparing the fold increase in P-p65 when treated with TNF-ɑ. Fold-change is relative to untreated condition. All experiments were performed in experimental triplicate (n=3) and standard error is shown in all graphs. Pair-wise t-tests were performed.

### MmuPV1 and HPV16 do not alter IKK complex protein abundance

We next aimed at determining the mechanism by MmuPV1 and/or HPV16 E6 impairs NF-κB signaling, specifically do they reduce IKK complex protein abundance. To address this, we performed immunoblot analysis using antibodies against IKKɑ, IKKβ, and IKKγ on protein extracts from the various cell lines used in our study. In mouse keratinocytes (MKs) that stably expressing WT MmuPV1 E6, MmuPV1 E6^R130A^, or vector control, we observed no change in the steady state abundance of IKKɑ, IKKβ, and IKKγ (**Fig. 7A & B**). We next analyzed MKs containing recircularized WT MmuPV1 or MmuPV1 E6^R130A^ genomes and dsRed control MKs. We found no significant change in the protein abundance of IKKɑ, IKKβ, and IKKγ (**Fig. 7C & D**). Finally, we examined IKK complex protein abundance comparing control, 16E6-expressing, 16E7-expressing, and 16E6E7- expressing N/TERT1 cells. We observed no significant change in the abundance of IKKɑ, IKKβ, and IKKγ in our N/TERT1 cells (**Fig. 7E & F**). We confirmed 16E6 and 16E7 expression by performing immunoblot analysis for p53 and pRB (**Fig. 7E**). Therefore, our data suggests that MmuPV1 and HPV16 E6 do no change IKK complex protein abundance as the mechanism by which they impair NF-κB signaling.

**Figure 7.**
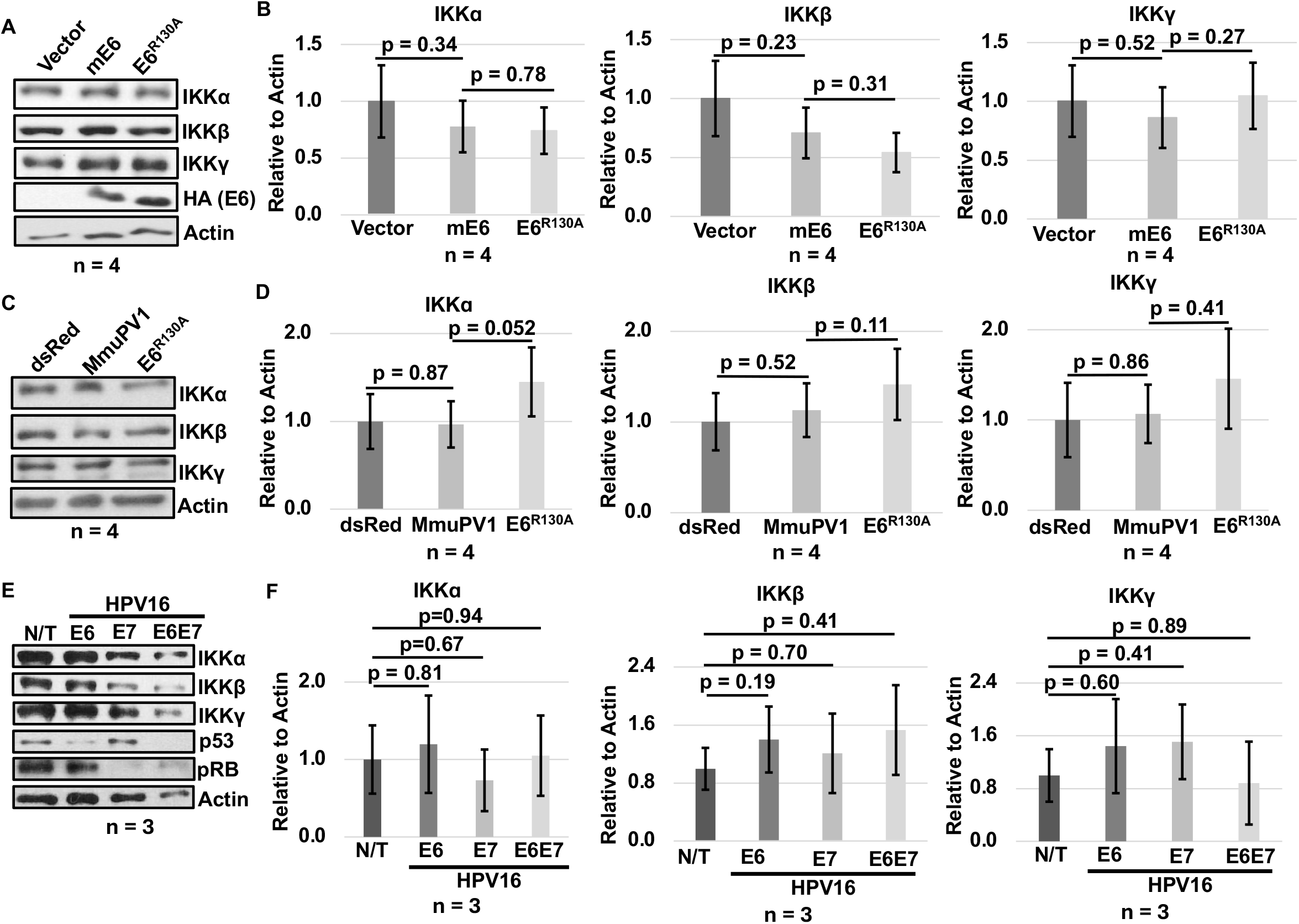
MmuPV1 and HPV16 E6 do not reduce IKK complex protein levels. (**A)** Immunoblot analysis was performed on extracts generated from mouse keratinocytes (MKs) stably expressing wild-type (WT) MmuPV1 E6 (mE6), MmuPV1 E6^R130A^, or vector control using antibodies against IKKα, IKKβ, IKKγ, HA (MmuPV1 E6), and actin. Representative blot is shown and analysis was performed in biological quadruplicate (n=4). (**B**) Quantification of immunoblot analysis shown in panel **A**. (**C)** Immunoblot analysis was performed on extracts generated from MKs containing WT MmuPV1 genome, MmuPV1 E6^R130A^ genome, or dsRed control using antibodies against IKKα, IKKβ, IKKγ, and actin. Representative blot is shown and analysis was performed in biological quadruplicate (n=4). **(D**) Quantification of immunoblot analysis shown in panel **C**. (**E)** Immunoblot analysis was performed on extracts generated from N/TERT1 cells expressing HPV16 E6, 16 E7, 16 E6/E7, or control cells using antibodies against IKKα, IKKβ, IKKγ, p53 (16 E6 expression control), pRB (16 E7 expression control), and actin. Representative blot is shown and analysis was performed in experimental triplicate (n=3). **(F**) Quantification of immunoblot analysis shown in panel E. Standard error is shown in all graphs.

### MmuPV1 E6 increases nuclear IKKɑ localization

In a recent study, IKKɑ has been shown to be both a cytoplasmic and nuclear protein in mouse keratinocytes (51). The ratio of nuclear to cytoplasmic IKKɑ has been reported to shift IKKɑ’s activities in keratinocytes (51). Importantly, increased nuclear IKKɑ has been reported to reduce NF-κB signaling (51). To determine whether MmuPV1 E6 alters IKKɑ localization as the mechanism by which MmuPV1 E6 impairs NF-κB signaling, we performed cellular fractionation analysis using the Rapid, Efficient, and Practical (REAP) method followed by immunoblot analysis (70). We found that MmuPV1 E6-expressing mouse keratinocytes (MKs) have significantly increased nuclear localization of IKKɑ compared to vector control MKs (**Fig. 8A & B**). The MmuPV1 E6^R130A^ mutant showed localization pattern consistent with vector control MKs and was significantly lower than WT MmuPV1 E6 (**Fig. 8A & B**). We also found that MmuPV1 E6 and E6^R130A^ are both a nuclear and cytosolic protein as expected with no significant difference in localization (**Fig. 8A & C**) (71). We used Lamin B1 and β-tubulin as controls for the fractionation (**Fig. 8A**). To determine whether MmuPV1 increases the nuclear localization of IKKɑ in the context of the intact viral genome, we repeated the fractionation assay in MKs containing recircularized WT MmuPV1 or MmuPV1 E6^R130A^ genomes and compared them to dsRed control MKs. The WT MmuPV1 genome-containing MKs showed a significant increase in nuclear IKKɑ compared to the dsRed control MKs (**Fig. 8D & E**). The MmuPV1 E6^R130A^ genome containing cells had a localization pattern similar to the dsRed control MKs and significantly lower nuclear IKKɑ compared to WT MmuPV1 genome containing cells (**Fig. 8D & E**). Lamin B1 and β-Tubulin were enriched in their respective compartments (**Fig. 8D**). These results suggest to us that MmuPV1 E6 increases nuclear IKKɑ which is dependent on binding since the MmuPV1 E6^R130A^ mutant does not interact with IKKɑ. This is also the first observation of a papillomavirus E6 oncoprotein altering IKKɑ localization.

**Figure 8.**
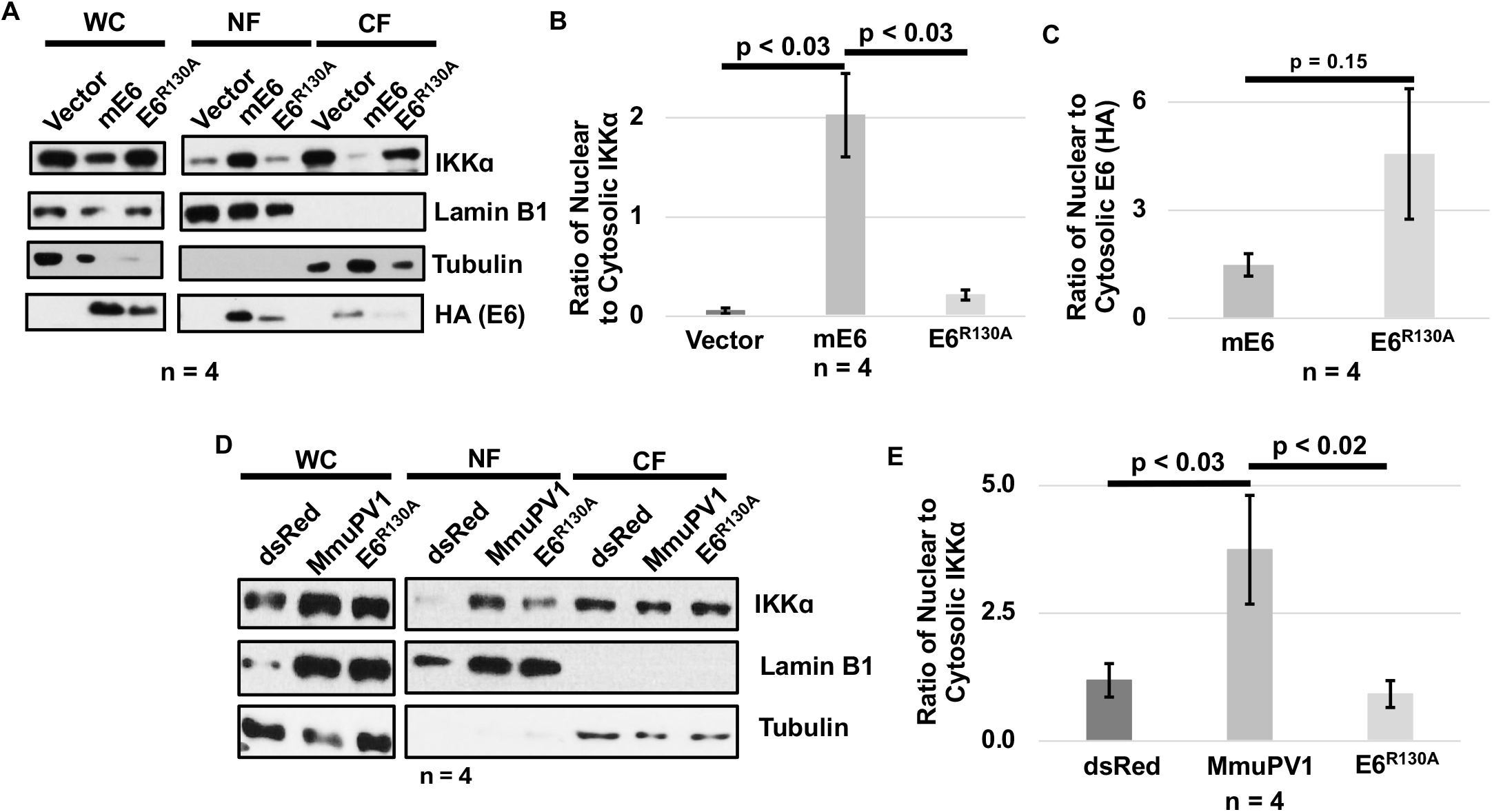
MmuPV1 promotes increased nuclear accumulation of IKKɑ which is binding dependent. **(A)** Mouse keratinocytes (MKs) stably expressing vector, wild-type (WT) MmuPV1 E6 (mE6), or MmuPV1 E6^R130A^ were subjected to cellular fractionation using the REAP method. Immunoblot analysis was performed on whole-cell (WC), nuclear (NF), and cytosolic (CF) fractions using antibodies against IKKɑ, Lamin B1 (nuclear fraction control), Tubulin (cytosolic fraction control), and HA (MmuPV1 E6). Representative blot is shown. **(B)** Quantification of the nuclear-to-cytosolic IKKα ratio of immunoblot shown in panel **A**. Standard error is shown. Pairwise t-test was performed. **(C)** Quantification of the nuclear-to-cytosolic ratio of MmuPV1 E6 was performed on immunoblot analysis shown in panel **A**. Standard error is shown. Pairwise t-test was performed. **(D)** MKs maintaining the intact WT MmuPV1genome, MmuPV1 E6^R130A^ genomes, or dsRed control were subjected fractionation with the REAP method. WC, NF, and CF fractions were subjected to immunoblot analysis using antibodies against IKKα, Lamin B1, and β-tubulin. Representative blot is shown. **(E)** Quantification of the nuclear-to-cytosolic IKKα ratio was performed on immunoblot analysis shown in panel **D**. Standard error is shown and pair-wise t- test was performed. All experiments were performed in biological quadruplicate (n=4).

### HPV16 E6 increases nuclear IKKɑ but not through a direct interaction with IKKɑ

Considering HPV16 E6 disrupts NF-κB signaling in human keratinocytes, we wanted to determine whether HPV16 E6 (16E6) promotes nuclear localization of IKKɑ. We performed fractionation analysis using the Rapid, Efficient, and Practical (REAP) method on control, 16E6-expressing, 16E7-expressing (negative control), and 16E6E7-expressing N/TERT1 cells. We found that 16E6 like MmuPV1 E6 significantly increases the nuclear localization of IKKɑ (**Fig. 9A & B**). Both Lamin B1 and β-Tubulin showed the anticipated enrichment in our fractionation analysis (**Fig. 9A**). To determine whether this increase localization was due to an interaction between 16E6 and IKKɑ, we performed Co-IP/WB analysis on control, HA-tagged 16E6, and HA-tagged 16E7 N/TERT1 cells. 16E6 and 16E7 are HA-tagged in our N/TERT1 cells. We did not observe an interaction between 16E6 or 16E7 and the IKK complex proteins (**Fig. 9C**). We used E6AP and pRB as controls for Co-IP/WB analysis (**Fig. 9C**). Our data suggests to us that HPV16 E6 alters IKKɑ localization through indirect mechanisms instead of through a direct interaction like MmuPV1 E6.

**Figure 9.**
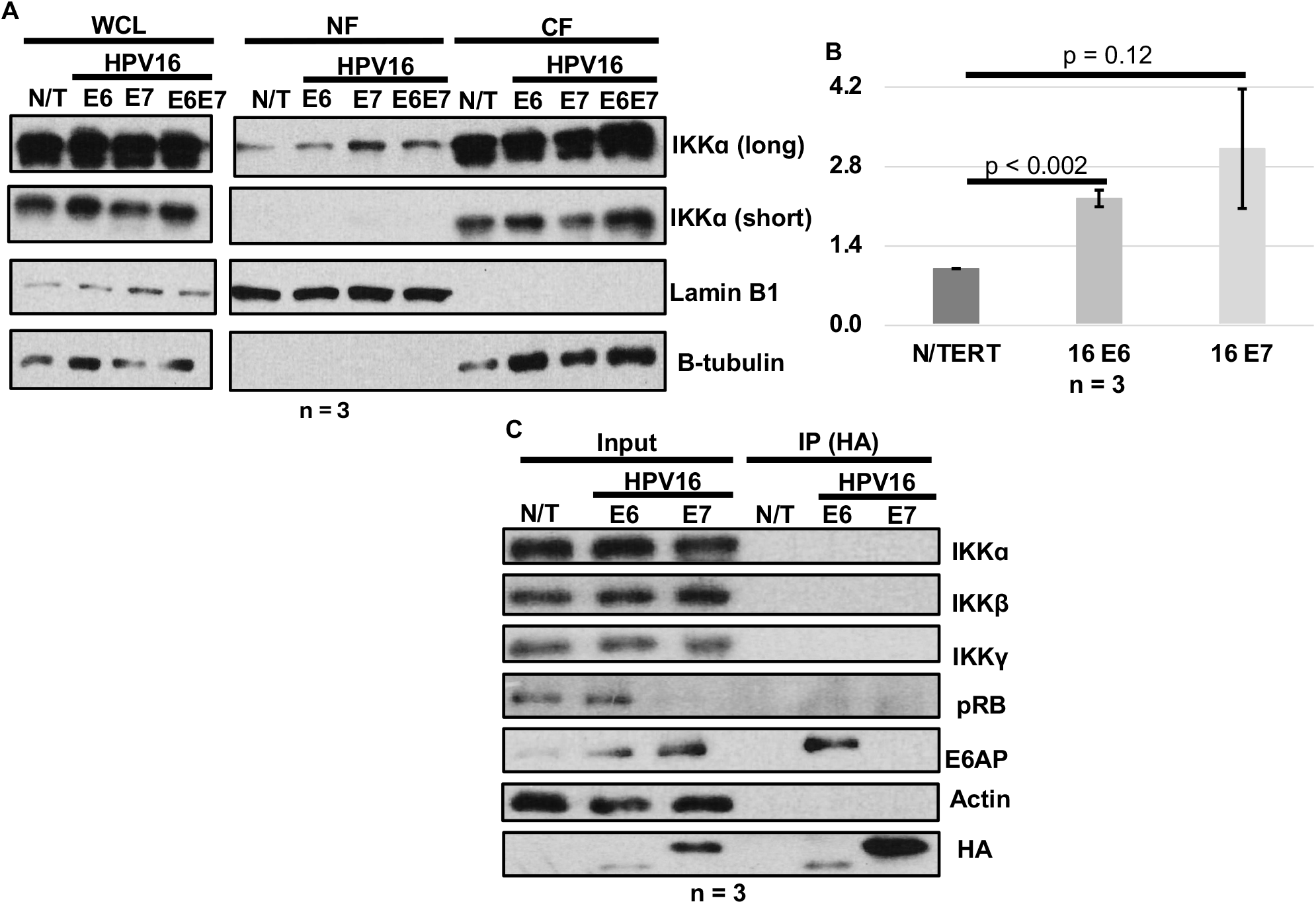
HPV16 E6 promotes increased accumulation of IKKɑ through an indirect mechanism. (**A**) Control N/TERT-1 cells and N/TERT1 cells stably expressing HPV16 E6, E7, or E6/E7 were separated into whole-cell lysate (WCL), nuclear (NF), and cytosolic (CF) fractions. Immunoblot analysis was performed on WCL, NF, and CF using antibodies against IKKɑ, LaminB1 (nuclear fraction control), and Tubulin (cytosolic control). Representative blot is shown. (**B**) Quantification of the nuclear-to-cytosolic IKKα ratio of immunoblot analysis shown in panel **A**. Pair-wise t-test was performed. (**C**) Lysates from parental N/TERT-1 cells and cells expressing HA-tagged HPV16 E6 or E7 were subjected to Co-IP/WB analysis using anti-HA antibodies. Immunoblot analysis was performed using antibodies against IKKɑ, IKKβ, IKKγ, pRB (16 E7 control), E6AP (16 E6 control), Actin, and HA (detects both 16 E7 and 16E7) on 10 μg inputs and immunoprecipitants. Representative image is shown. All studies were performed in experimental triplicate (n=3).

### MmuPV1 E6 promotes transcriptional changes associated with nuclear IKKɑ

A recent publication showed that increased nuclear IKKɑ can promote several transcriptional changes in keratinocytes (66). Considering that MmuPV1 E6 promotes a strong change in nuclear localization, we wanted to determine if transcriptional changes in our RNAseq data could correlate with increased nuclear IKKɑ. Using Gene Set Enrichment Analysis (GSEA), we probed our published RNAseq analysis comparing MmuPV1 E6-expressing mouse keratinocytes (MKs) to vector control MKs specifically looking at the Hallmark gene sets which are curated by the Broad Institute (72). Consistent with the published study, MmuPV1 E6 expression in mouse keratinocytes promotes a significant positive enrichment in the HALLMARK_MYC_Targets_V1 gene set (**Fig. 10A**). Additionally, MmuPV1 E6 promotes a significant negative enrichment in the HALLMARK_FATTY_ACID_METABOLISM (**Fig. 10B**), HALLMARK_ADIPOGENESIS (**Fig. 10C**), HALLMARK_INTERFERON_ALPHA_RESPONSE (**Fig. 10D**), HALLMARK_INTERFERON_GAMMA_RESPONSE (**Fig. 10E**), and HALLMARK_IL6_JAK_STAT3_SIGNALING (**Fig. 10F**) gene sets. All our observations are consistent with the prior study (51). Therefore, our data suggest that the increase in nuclear IKKɑ correlates with several transcriptional changes promoted by MmuPV1 E6 that likely contribute to persistent infection.

**Figure 10.**
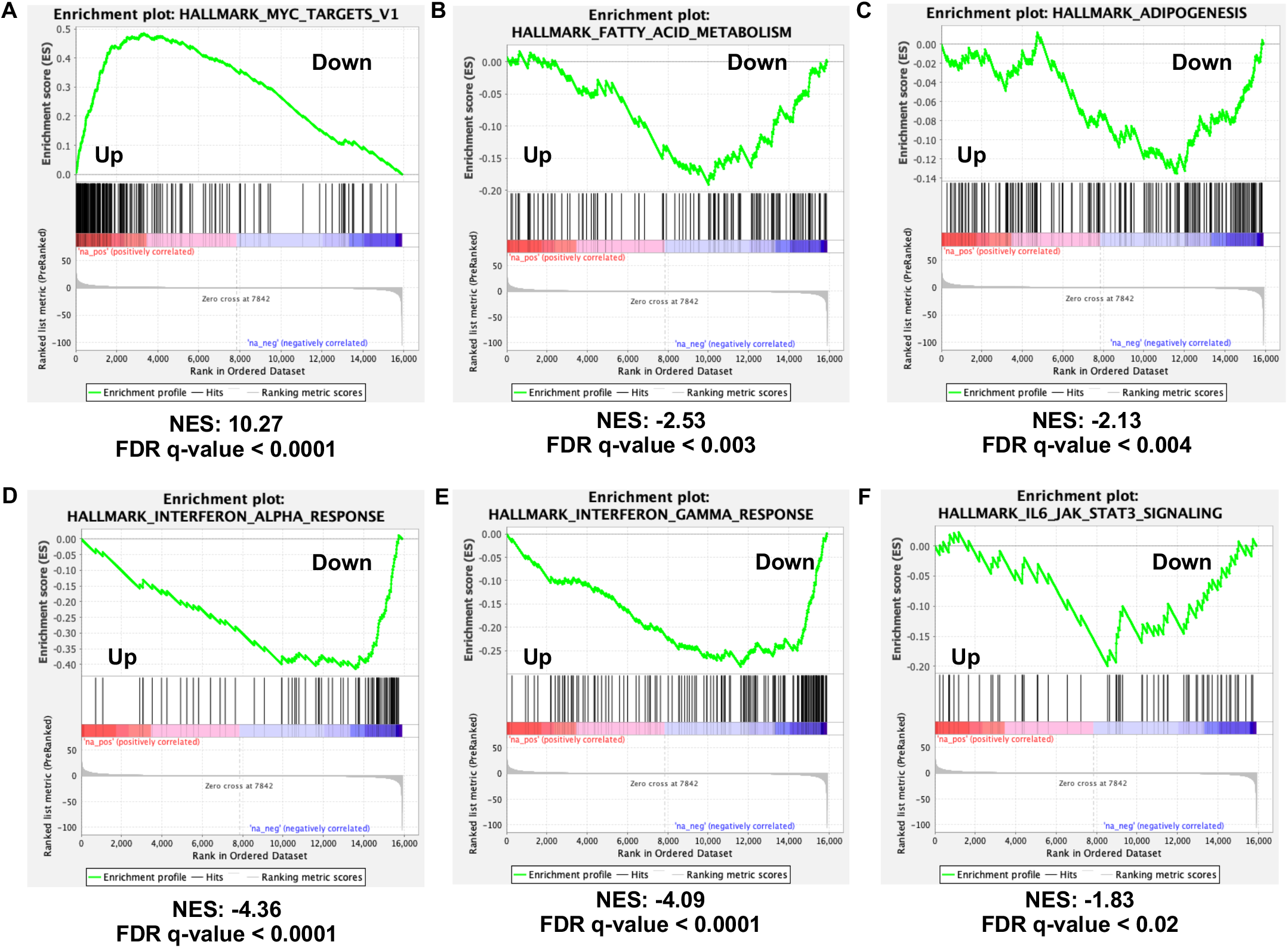
MmuPV1 E6 promotes transcriptional changes that are associated with increased nuclear IKKɑ accumulation. Gene set enrichment analysis was performed ou our previously published RNA-seq data from mouse keratinocytes stably expressing MmuPV1 E6 or vector control mouse keratinocytes (MKs) (66). Normalized enrichment score (NES) and false- discovery rate (FRD) q-values are shown. The six Hallmark gene sets shown were selected based on their reported enrichment in mouse skin expressing nuclear-targeted IKKɑ (51). The Hallmark_MYC_Targets_V1 **(A)** gene set was positively enriched in MmuPV1 E6-expressing MKs. The Hallmark_Fatty_Acid_Metabolism **(B)**, Hallmark_Adipogenesis **(C)**, Hallmark_Interferon_Alpha_Response**(D)**, Hallmark_Interferon_Gamma_Response **(E)**, and Hallmark_IL6_JAK_STAT3_Signaling **(F)** were negatively enriched in MmuPV1-E6 expressing MKs.

## Discussion

Our collective data in this study has identified novel interacting partners for a papillomavirus E6 oncoprotein – IKKɑ, IKKβ, and IKKγ/NEMO (**Fig 3**). The functional consequence of that interaction is MmuPV1 E6 impairs NF-κB signaling in mouse keratinocytes (**Figs. 1**, **4-5**). Importantly, we found at HPV16 E6 also impairs NF-κB signaling in human keratinocytes (**Fig 6**). This observation is key as this is the first functional similarity between MmuPV1 E6 and a high-risk HPV E6. To date, MmuPV1 E6 is considered more genetically related to cancer-associated cutaneous HPV E6 oncoproteins (24–26, 65, 73). This observation is based upon that MmuPV1 E6 interacts with MAML1 and SMAD2/3 which are targets for cutaneous HPV E6 (65). However, MmuPV1 E6 also interacts with IRF3 like high-risk HPV E6 (27, 74, 75). Cutaneous HPV E6 does impair IRF3 activity but does not directly interact with IRF3 (74). Therefore, our collective data is beginning to suggest that MmuPV1 E6 may not fall into the clear separation between high-risk mucosal and cutaneous HPVs as previously thought. More studies are needed to better define the shared biology and interacting partners between MmuPV1 E6 and HPV 16 including high-risk and cutaneous.

The MmuPV1 E6^R130A^ mutant was defective in binding with IKK complex proteins and could no longer impair NF-κB signaling (**Figs 3-5**). These results suggest that the E6^R130A^ mutant is a multifunctional mutant and impacts a number of MmuPV1 E6 activities. The MmuPV1 E6^R130A^ mutant is hypothesized to be an LXXLL-motif binding mutant due to the binding defect with the LXXLL-motif containing protein MAML1 (65). However, IKKɑ, IKKβ, and IKKγ are not known LXXLL-motif containing proteins. After examining IKKɑ, IKKβ, and IKKγ for LXXLL motifs, we only found 1 in IKKɑ. However, the potential LXXLL motif in IKKɑ does not contain any acidic residues in the XX region of the motif like the other well described LXXLL-motif containing host proteins, i.e., MAML1 and E6AP each contain at least 1 acidic residue (24). However, our work does not go into defining whether the potential LXXLL-motif in IKKɑ contributes to the interaction between IKKɑ and MmuPV1 using IKKɑ mutants. Future studies could aim to determine whether MmuPV1 E6 interacts with IKKɑ through an LXXLL-motif.

Many previous studies have shown that high-risk HPV E6 and E7 target NF-κB signaling. The initial study that showed high-risk HPV E7 targets NF-κB signaling in cervical transformation zone cells (37, 38). Interestingly, they found overexpressing p65 or expression of a dominant negative IκBɑ mutant that lacks phosphorylation sites impacted the growth of cervical cells (38). An interpretation of this study is that NF-κB signaling is anti-proliferative in normal cervical cells and impairs some of the early stages of HPV-associated disease. A later study showed HPV16 E7 interacts with the IKK complex proteins to attenuate NF-κB signaling (37). We did not see similar results in our N/TERT1 cell studies (**Fig. 6**). It should be noted that this study used rat cells and U2OS cells instead of keratinocytes in their study (37). However, it does not remove the possibility that our experimental design fails to capture every aspect of HPV16 E7. Another study on high-risk HPVs impact on NF-κB showed that high-risk HPV increase UCHL1 to degrade IKKγ (NEMO) to inhibit NF-κB signaling (36). In future studies, we could mine our previously published RNAseq data sets for MmuPV1 E6 and E7 to determine their impact on UCHL1 (66, 76). In contrast, there are studies that show high-risk HPV E6 activates NF-κB signaling (28, 31, 32, 39, 41, 45). HPV positive cervical cancers have increased expression of NF-κB responsive genes (31, 47). A number of mechanisms have been proposed including targeting of TLR molecules (32). However, this study used overexpression plasmids for E6 and TLR proteins in HEK293 cells which are not epithelial in origin (32). In our Co-IP/WB analysis, we used stably expressing human keratinocytes (**Fig. 9**). The differences in the model systems likely explain the differences in our observations. Another study showed that high-risk HPV E6 reduces the expression of p105 by degrading NFX1-91 and leads to an activation of NF-κB signaling (28). They were able to correlate the increase in NF-κB signaling with a reduction in the expression of genes from the HPV early and late promoters (28). High-risk HPV E6 also increases the expression of cIAP-2 through NF- κB to protect the cells from apoptosis (39). This complex interaction of high-risk HPV E6 with NF- κB signaling may reflect the pro- and anti-pathogenic roles of NF-κB in HPV-associated disease. Therefore, more studies are needed to better understand how HPV and MmuPV1 E6 modulate this signaling cascade to promote disease and cancer development.

The impact of high-risk HPV16 E6 was most apparent in our co-culture of N/TERT1 cells with J2 3T3 fibroblasts (**Fig. 6**). These results are interesting because recent studies have found that feeders can significantly impact the biology of keratinocytes and the behavior of high-risk HPV E6 and E7 (77–79). Feeders elevate expression of proliferation genes, promote increased and decreased transcription of various chemokines and cytokines, and the innate immune response (77). Interestingly, feeders promote maintenance of episomal HPV genomes in keratinocytes (79). Feeders also contribute to the response of keratinocytes to drug treatment (78). Therefore, we are not surprised by these results in our N/TERT1 study showing feeders may contribute to HPV16 E6’s repression of TNFɑ activation of NF-κB. However, the impact of feeders on experimental design should be considered as we continue our work on HPV and MmuPV1. The exact contribution of feeders and what feeders do to support these changes remains an unanswered question in the field and will require more studies to better understand how they contribute to papillomavirus persistent infection and disease development.

Our most interesting finding is that both MmuPV1 E6 and HPV16 E6 promote nuclear accumulation of IKKɑ (**Figs. 8 & 9**). Nuclear IKKɑ has been reported in Hepatitis B infection but never during a papillomavirus infection (80, 81). In the case of the skin, IKKɑ plays a critical role in skin development and is a tumor suppressor gene (48, 49, 51–54). Mice that have constitutive knockout of IKKɑ do not develop normal skin morphology (48, 51, 53). IKKɑ also contributes to keratinocyte differentiation (48, 49, 53, 54). Therefore, IKKɑ plays a very similar role as MAML1/NOTCH signaling in keratinocyte biology (82–89). This makes our finding that the MmuPV1 E6^R130A^ mutant having a defect in binding to both IKKɑ and MAML1 interesting. The prior work on the E6^R130A^ mutant has found it does not impair differentiation like WT MmuPV1 E6 (65, 67). Our results make the role of MAML1 and IKKɑ less clear as it is possible that loss of binding both is why the MmuPV1 E6^R130A^ mutant cannot impair differentiation. Additionally, we may not be accounting for other interacting partners that require amino acid R130 in MmuPV1 E6 that could also be contributing to this phenotype. Therefore, future studies are needed to better define the individual contributions of various interacting partners of MmuPV1 E6.

Nuclear IKKɑ has been linked to a number of transcriptional changes observed in mouse skin (51). Importantly, we observe similar phenotypes in our RNAseq analysis comparing MmuPV1 E6 to vector control mouse keratinocytes (51) (**Fig. 10**). An interesting result is the increase in c-Myc responsive gene expression (**Fig. 10A**). High-risk HPV E6 has previously been reported to increase c-Myc activity (30, 90–94). Nuclear IKKɑ has been shown to stabilize active c-Myc from degradation (95). Therefore, MmuPV1 E6’s increase in nuclear IKKɑ might be promoting c-Myc activity which likely contributes to persistent infection and cancer development. The impact on fatty acid metabolism is particularly interesting as MmuPV1 infection alters lipid metabolism (96, 97). MmuPV1 E6 has not been directly linked to these changes and remains an interesting and unanswered question. The impact on innate immune signaling is not unexpected as NF-κB plays a key role in the innate immune response (13) (**Fig 10**). However, we cannot rule out MmuPV1 E6’s interaction with IRF3 which also impact the innate immune response (27, 98, 99). The only unexpected result is the reduction in IL-6 and STAT3 signaling. STAT3 has very clearly been shown to be increased by high-risk HPV E6 and plays a role in viral replication and proliferation of HPV+ keratinocytes (100). IL-6 has also been shown to be enriched in HPV+ cancer (101–103) and promotes growth of keratinocytes and wound healing (104, 105). We do observe a baseline reduction in IL-6 expression that likely explains these results (**Fig 1**). It is important to note that the HALLMARK_IL6_JAK_STAT3_SIGNALING gene set has the weakest negative enrichment which could reflect a more nuanced impact on this signaling cascade by MmuPV1 E6. Future studies are needed to address all these observations.

To conclude, we have found that MmuPV1 E6 interacts with the IKK complex proteins which is the first time this has been observed in keratinocytes. This interaction contributes to MmuVP1 E6’s ability to impair NF-κB signaling in keratinocytes. The impairment in NF-κB is shared with high-risk HPV16 E6 in human keratinocytes. Our most exciting result is that MmuPV1 and HPV16 E6 increase nuclear IKKɑ localization. Consistent with this observation, MmuPV1 E6 promotes several transcriptional changes that correlate with increased nuclear IKKɑ and these changes likely contribute to persistent infection. Future work in better identifying how nuclear IKKɑ promotes these changes during MmuPV1 infection are needed to address these exciting observations.

## Materials and Methods

### Cell Culture

NIH J2 3T3 fibroblasts obtained from ATCC were maintained in Dulbecco’s modified Eagle medium (DMEM; Gibco) supplemented with 10% bovine calf serum (Cytiva) and penicillin-streptomycin (Gibco). 293FT cells obtained from ATCC were maintained in DMEM supplemented with 10% fetal bovine serum, penicillin-streptomycin, and 200 μg/mL G418 (Geneticin; Gibco). HCT116 cells obtained from ATCC were maintained in McCoy’s 5A medium (Gibco) supplemented with 10% fetal bovine serum (Cytiva) and penicillin-streptomycin. Mouse keratinocytes (MKs) were isolated from neonatal FVB/N mouse skin as previously described and cultured in F-medium containing 10 μM ROCK inhibitor Y-27632 on mitomycin C (MMC)-treated J2 3T3 feeder cells (106). J2 3T3 fibroblasts were treated with MMC for 2 hrs before plating in F- medium with or without Y-27632, as required for the indicated assay. Early-passage MKs were transduced in keratinocyte serum-free medium (KSFM; Gibco) with retroviral or lentiviral vectors encoding MmuPV1 E6, MmuPV1 E6^R130A^, or the corresponding vector control (65, 66, 76). At 48 hrs post-transduction, transduced cells were selected with the appropriate selection agent, i.e., 200 μg/mL G418 (Gibco) for retrovirus or 7 μg/mL blasticidin S hydrochloride (Gibco, cat. no. A11139-03) for lentivirus and maintained in F-medium containing Rock inhibitor containing the appropriate selection agent. N/TERT-1 human keratinocytes and derivatives expressing HPV16 E6, HPV16 E7, or HPV16 E6/E7 were a gift from Drs. Claire D. James and Iain M. Morgan (Virginia Commonwealth University) and were described previously (77, 94).

### Plasmids

All MmuPV1 E6, MmuPV1 E6^R130A^, and control plasmids – pLentiN, pLXSN, and pN-CMV – were a gift from Dr. Karl Munger (Tufts) (65). pcDNA6-dsRed and plasmids containing full-length wild-type or E6^R130A^ MmuPV1 genomes were a gift from Dr. Paul Lambert (University of Wisconsin-Madison) (68). Full-length viral genomes were released from their bacterial plasmid backbones with XbaI and recircularized by intramolecular ligation with T4 DNA ligase as described previously (61, 62, 68).

### Lentivirus and Retrovirus Production

MmuPV1 E6, MmuPV1 E6^R130A^, and vector-control lentiviral and retroviral particles were generated as described previously (66, 76). For lentivirus production, pLenti-N constructs were co-transfected with psPAX2 and pMD2.G packaging plasmids (Addgene). For retrovirus production, pLXSN constructs were co-transfected with pCL- 10A1 (Fisher Scientific). Transfections were performed with Lipofectamine 2000 (Invitrogen, 11668-019) according to the manufacturer’s instructions.

### Cell Counting Assays

Short- and long-term growth analyses were performed as previously described (66, 76). All growth assays were performed in F-medium without Y-27632.

### TNFα Stimulation

MKs were seeded at 5X10⁴ cells per well in two wells of a six-well plate containing MMC-treated J2 3T3 feeder cells in F-medium supplemented with Y-27632. N/TERT- 1 cells were seeded at 1X10⁵ cells per well in two wells of a six-well plate containing feeder cells in F-medium without Y-27632. The following morning, all cultures were switched to F-medium without Y-27632. After 24 hrs, feeder cells were removed, cultures were washed twice with DPBS, and 2 mL prewarmed F-medium without Y-27632 was added per well. Cells were stimulated with recombinant mouse TNFɑ (Sigma-Aldrich, cat. no. T7539) at 10 ng/mL for 10 min, with matched unstimulated cultures processed in parallel. Cells were rapidly washed twice with ice-cold DPBS and lysed in RIPA buffer supplemented with cOmplete Mini EDTA-free protease inhibitor cocktail (Roche, cat. no. 11836170001) and phosphatase inhibitor cocktails II and III (Sigma-Aldrich, P5726 and P0044).

### RNA Isolation and Quantitative RT-PCR

Total RNA was extracted from TRIzol lysates by phenol-chloroform extraction according to the manufacturer’s protocol. cDNA was synthesized using the High-Capacity cDNA Reverse Transcription Kit (Applied Biosystems) according to the manufacturer’s instructions. Quantitative PCR was performed with SYBR Green chemistry (Bio-Rad) on a CFX96 real-time PCR detection system (Bio-Rad). Primer sequences are listed in Table 1 (66, 68, 76).

**Table 1.**
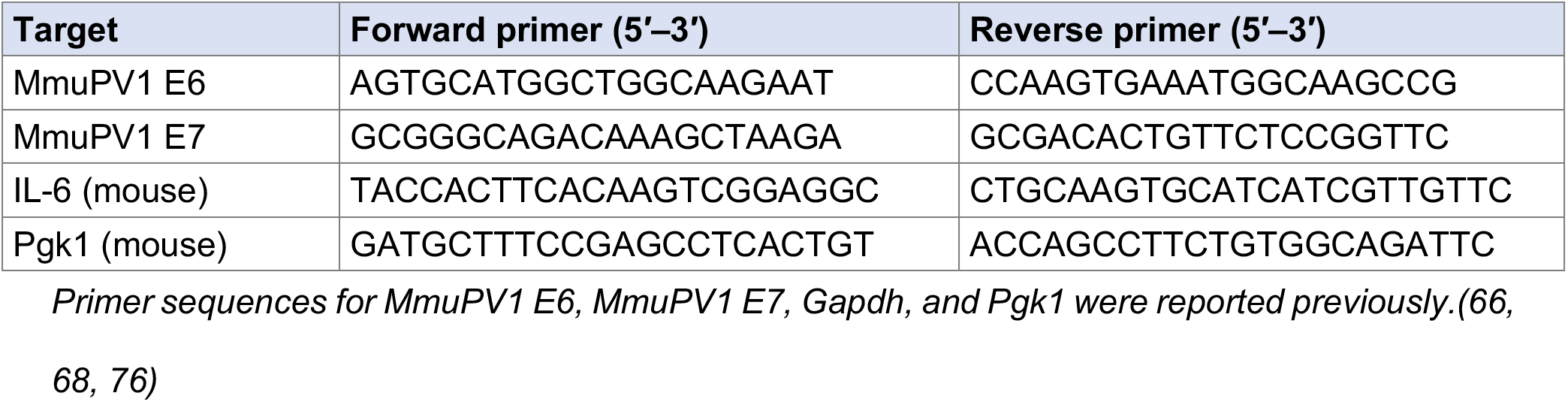
Oligonucleotides used for quantitative RT-PCR.

### Fractionation Assay

Subcellular fractionation was performed using the Rapid, Efficient and Practical (REAP) method (70). Near confluent MK and N/TERT1 cultures were used. At near confluence, feeder cells were removed and washed twice with ice-cold DPBS. Cells were scraped into 1 mL ice-cold PBS and pelleted. Pellets were resuspended in 450–900 μL ice-cold PBS containing 0.1% NP-40 based on pellet size. 150–300 μL was reserved as the whole-cell fraction. The remaining lysate was centrifuged at maximum speed for 10 s, and 150–300 μL supernatant was collected as the cytosolic fraction. The nuclear pellet was washed once with 1 mL PBS containing 0.1% NP-40, centrifuged at maximum speed for 10 s, and resuspended in 180 μL 1X SDS sample buffer. Whole-cell and nuclear fractions were sonicated twice for 5 s at 50% amplitude and analyzed by immunoblotting.

### Immunoblot Analysis

For steady-state protein analysis, sub-confluent keratinocyte cultures were lysed in RIPA buffer as described above. Protein concentration was determined using the Pierce Dilution-Free Rapid Gold BCA Protein Assay Kit (Thermo Fisher Scientific) according to the manufacturer’s instructions. Immunoblot analysis was performed as previously described (76). Amersham Protran nitrocellulose membranes (Cytiva) were used - 0.2-μm membranes (cat. no. 10600006) for 15% gels and 0.45-μm membranes (cat. no. 10600007) for 10% gels. Membranes were blocked in 5% nonfat milk or 5% BSA in PBS-T at room temp (RT), based upon primary antibody, and incubated with primary antibodies at 4°C overnight. All antibodies are listed in Table 2. The next day after washing, membranes were incubated with the appropriate HRP- conjugated secondary antibody (Rockland Immunochemicals) in 5% nonfat milk in PBS-T for 1 hr at RT. Blots were developed using Pierce ECL Western Blotting Substrate or SuperSignal West Pico PLUS Chemiluminescent Substrate (Thermo Fisher Scientific) according to the manufacturer’s instructions. Band intensities were quantified using ImageJ (107, 108).

**Table 2.** Antibodies used in this study.

| **Target/reagent** | **Host/clone** | **Supplier and catalog no.** | **Dilution** |
|---|---|---|---|
| Phospho-NF-κB p65 (Ser536) | Rabbit mAb, 93H1 | Cell Signaling Technology, #3033 | WB; 1:1000 |
| NF-κB p65 | Rabbit mAb, D14E12 | Cell Signaling Technology, #8242 | WB: 1:1000 |
| NF-κB1 p105/p50 | Rabbit mAb, D4P4D | Cell Signaling Technology, #13586 | WB; 1:1000 |
| NF-κB1 | Rabbit mAb, D7H5M | Cell Signaling Technology, #12540 | IIHC 1:1000 |
| NF-κB2 p100/p52 | Rabbit polyclonal | Cell Signaling Technology, #4882 | WB; 1:1000 |
| IKKα | Rabbit polyclonal | Cell Signaling Technology, #2682 | WB; 1:1000 |
| IKKβ | Rabbit mAb, D30C6 | Cell Signaling Technology, #8943 | WB; 1:1000 |
| IKKγ (NEMO) | Rabbit polyclonal | Cell Signaling Technology, #2685 | WB; 1:1000 |
| MAML1 | Rabbit mAb, D3K7B | Cell Signaling Technology, #12166 | WB; 1:1000 |
| IRF3 | Rabbit mAb, D83B9 | Cell Signaling Technology, #4302 | WB; 1:1000 |
| SMAD2/3 | Rabbit mAb, D7G7 | Cell Signaling Technology, #8685 | WB; 1:1000 |
| UBE3A (E6AP) | Rabbit mAb, D10D3 | Cell Signaling Technology, #7526 | WB; 1:1000 |
| HA | Rabbit mAb, C29F4 | Cell Signaling Technology, #3724 | WB; 1:2000 |
| GFP | Rabbit polyclonal | Cell Signaling Technology, #2555 | WB; 1:1000 |
| p53 | Mouse mAb, 1C12 | Cell Signaling Technology, #2524 | WB; 1:1000 |
| pRb | Mouse mAb, 4H1 | Cell Signaling Technology, #9309 | WB; 1:1000 |
| β-actin | Rabbit polyclonal | Cell Signaling Technology, #4967 | WB; 1:1000 |
| Lamin B1 | Rabbit polyclonal | Cell Signaling Technology, #9087 | WB; 1:1000 |
| β-tubulin | Rabbit polyclonal | Cell Signaling Technology, #2146 | WB; 1:1000 |

### Co-Immunoprecipitation (Co-IP) Analysis

HCT116 cells were transiently transfected with 2 μg pN-CMV, pN-CMV HA-tagged MmuPV1 E6, or pN-CMV HA-tagged MmuPV1 E6^R130A^ using Lipofectamine 3000 (Invitrogen) and harvested 48 hrs after transfection. Stable-expression experiments used near confluent vector control, HA-tagged MmuPV1 E6, or HA-tagged MmuPV1 E6^R130A^ MKs or vector control, HPV16 E6, or HPV16 E7 N/TERT1s, as indicated. At the time of collection, feeders were removed, cells were washed with PBS, scraped, and pelleted at 500xg for 5 min. Cell pellets were lysed in EBC buffer supplemented with cOmplete Mini EDTA-free protease inhibitor cocktail and phosphatase inhibitor cocktails II and III as previously described (65). Lysates were rotated for 30 min at 4°C and clarified by centrifugation at 13,000 rpm for 20 min at 4°C. Following protein quantification, a 10 μg input aliquot was reserved per gel from each sample. At least 0.1 mg total protein was used for keratinocytes and 1 mg of protein for HCT116 cells. These amounts were incubated with 20 μL anti-HA agarose slurry (Sigma-Aldrich, A2095) for 1 hr at 4°C with rotation. Bound proteins were eluted in SDS sample buffer containing β- Mercaptoethanol. Input lysates and HA immunoprecipitants were analyzed by immunoblotting as described above.

### MmuPV1 Infection and Tissue Collection

Immunohistochemistry was performed on banked skin tissues collected in a previous study (68). The banked tissues were generated under a University of Wisconsin Institutional Animal Care and Use Committee-approved protocol (M005871).

### Immunohistochemistry (IHC) Analysis

IHC was performed as previously described (68). Briefly, 5 μm tissue sections were deparaffinized in xylene and rehydrated through a graded ethanol series. Antigen retrieval was performed in citrate buffer (pH 6.0). Sections were blocked for 1 hr at RT with 5% normal goat serum and incubated overnight at 4°C with primary antibody (Table 2). The following day, sections were incubated with a secondary antibody for 1 hr at room temperature and developed with DAB (Vector Laboratories, cat. no. SK-4100). Sections were counterstained with hematoxylin (Vector Labs, H-3404-100). Slides were dehydrated through graded ethanol, cleared in xylene, and coverslips mounted with Epredia Signature Series Mounting Medium (supplier no. 4112). Slides were imaged using a Zeiss Axio Imager.M2 microscope (Carl Zeiss Microscopy).

## Data Availability

All data is available in public databases. The RNA-seq data reported in this paper are available in the BioProject database with the accession number <u>PRJNA1007707</u>.

## Acknowledgements

We want to acknowledge Liyan Zhang who completed the initial drafts of this manuscript and completed the studies shown in Figures 3-9. This study was supported by grants awarded from the US NIH (R01 CA228543, R35 CA210807, P01 CA022443, F32 CA254019, and P41 GM108538). Startup funds from the Virginia-Maryland College of Veterinary Medicine (VMCVM) and seed funds from the VMCVM and Edward Via College of Osteopathic Medicine (VCOM) were used to support experiments performed in this study.

